# Excitatory and inhibitory intracortical circuits for orientation and direction selectivity

**DOI:** 10.1101/556795

**Authors:** L. Federico Rossi, Kenneth D. Harris, Matteo Carandini

**Affiliations:** UCL Institute of Ophthalmology, University College London, UK; UCL Institute of Neurology, University College London, UK

## Abstract

The computations performed by a neuron arise from the functional properties of the circuits providing its synaptic inputs. A prime example of these computations is the selectivity of primary visual cortex (V1) for orientation and motion direction. V1 neurons in layer 2/3 (L2/3) receive input mostly from intracortical circuits^1^, which involve excitation^2-9^ and inhibition^10-12^. To understand how an L2/3 neuron achieves its selectivity, therefore, one must characterize the functional organization of both its excitatory and inhibitory presynaptic ensembles. Here we establish this organization, and show how it predicts orientation selectivity and reveals a new cortical circuit for direction selectivity. We identified the presynaptic partners of pyramidal neurons in mouse V1 through rabies monosynaptic tracing^1,13^, and imaged the functional properties of the postsynaptic neuron and of its presynaptic ensemble. Excitatory presynaptic neurons were predominantly tuned to the postsynaptic neuron’s preferred orientation. Excitation and inhibition described an inverted Mexican hat, with inhibitory presynaptic neurons densest near the postsynaptic neuron and excitatory ones distributed more distally. Excitation and inhibition also differed in laminar origin: inhibitory presynaptic neurons concentrated in L2/3 while excitatory ones dominated in L4. The distribution of excitatory neurons in visual space was coaxial with the postsynaptic neuron’s preferred orientation and lay upstream of the neuron’s preferred direction. Inhibitory presynaptic neurons, instead, clustered more symmetrically around the postsynaptic neuron and favoured locations downstream of its preferred direction. These results demonstrate that L2/3 neurons obtain orientation selectivity from co-tuned neurons in L4 and beyond, and enhance it by contrasting an elongated excitatory input with a concentric inhibitory input. Moreover, L2/3 neurons can obtain direction selectivity through visually offset^14^ excitation and inhibition. These circuit motifs resemble those seen in the thalamocortical pathway^15-20^ and in direction selective cells in the retina^21,22^, suggesting that they are canonical across brain regions.

---

A fundamental question in neuroscience is how the specialized responses of one neuron are determined by the interplay of its excitatory and inhibitory presynaptic inputs. In L2/3 of the primary visual cortex, neurons respond preferentially to oriented stimuli that move in a specific direction. These responses are driven by thousands of inputs from excitatory and inhibitory ensembles distributed across cortical layers^23^. Excitatory presynaptic ensembles tend to be co-tuned with the postsynaptic neuron^1-8^, but it is unclear whether such co-tuning applies to inputs from all layers^1^, and whether it explains the selectivity of individual postsynaptic neurons for orientation and direction^1,2,4,6,10^. Moreover, the functional organisation of inhibitory presynaptic ensembles remains unknown, and it is thus unclear whether and how inhibitory ensembles shape either form of selectivity^10-12,14,24-26^. While studies *in vitro* support a close balance of synaptic inhibition and excitation^12^ both horizontally^27-29^ and across laminae^28^, neuroanatomy predicts marked differences in the cortical distribution of excitatory and inhibitory presynaptic ensembles^23^, whose functional relevance remains unexplored.

To understand the functional basis for the selectivity of individual V1 neurons in L2/3, we established their presynaptic ensemble across layers^1^, while distinguishing excitatory^2-9^ from inhibitory^10-12^ presynaptic partners (Figure 1a-d). First we electroporated 3-5 pyramidal neurons in L2/3 with genes for the avian receptor TVA, the rabies glycoprotein oG and the red marker dsRed, in mice expressing the calcium indicator GCaMP6 in cortical excitatory neurons (Figure 1a, b, Supplementary Figure 1). Second, we recorded the visual responses of the electroporated neurons (Figure 1a, b), selected one based on visual selectivity, and photo-ablated any additional ones (Supplementary Figure 2). Third, we injected a modified rabies virus^13^ expressing dsRed, which could only infect the TVA-positive target neuron, which also contained the glycoprotein required to propagate to presynaptic partners (Figure 1a, c). Lastly, we used two-photon imaging for anatomical reconstructions and to record responses from 10,000-14,000 excitatory neurons, including the postsynaptic neuron, its presynaptic ensemble, and the surrounding population (Figure 1c). Thanks to these images, we could identify the borders between layers (Figure 1c) and classify presynaptic neurons as excitatory or inhibitory based on the co-localisation of rabies-dsRed2 and GCaMP6 in the soma (Figure 1d, e, Supplementary Figure 3). For each postsynaptic neuron (N=13), we traced on average 123±26 local V1 presynaptic partners (ranging from 42 to 334). Most of these inputs where excitatory (median 59%), but a substantial fraction was inhibitory (Supplementary Figure 3).

**Figure 1.**
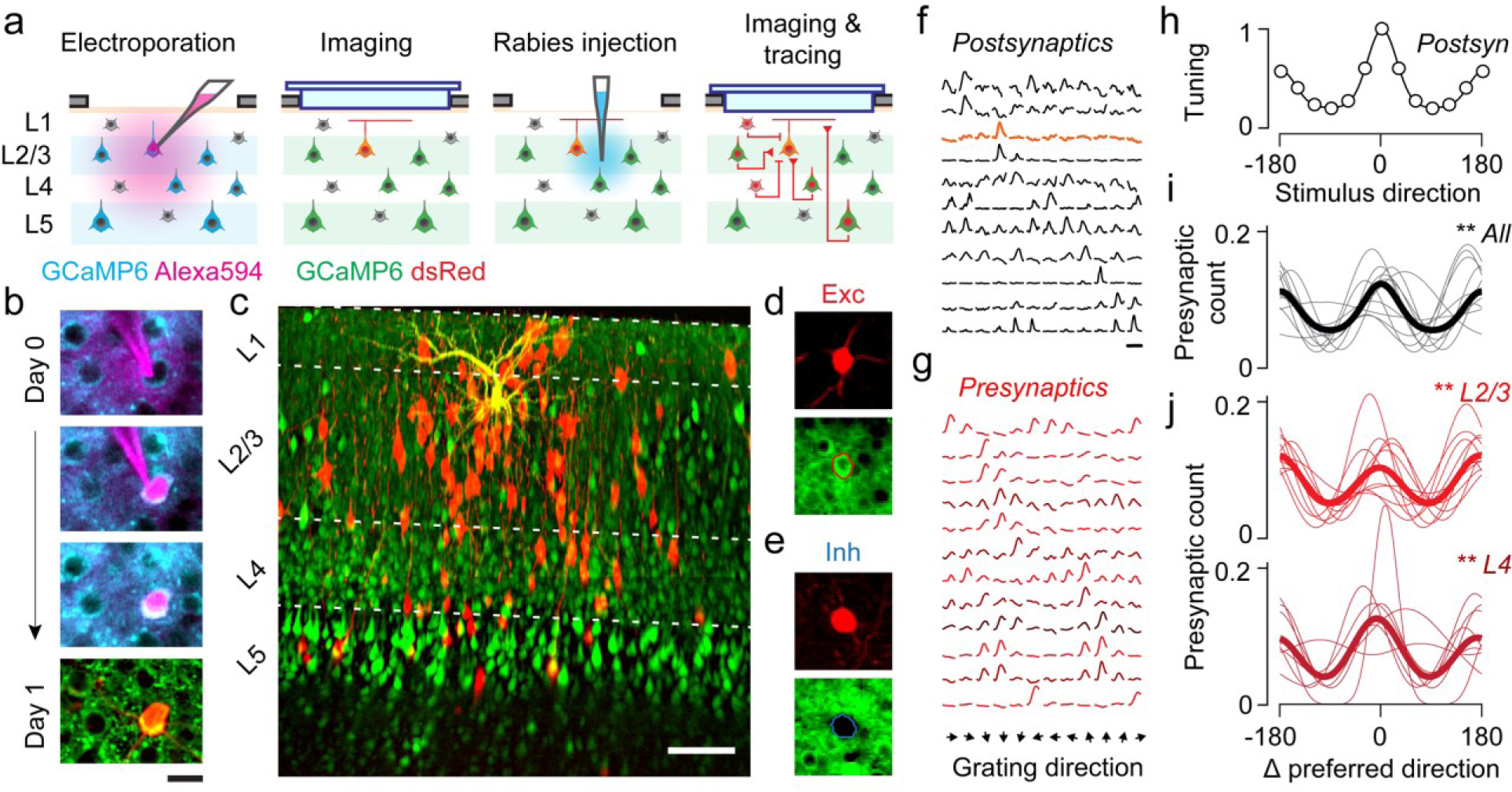
Tracing the excitatory and inhibitory neurons presynaptic to target L2/3 pyramidal neurons. (**a**) Experimental pipeline: electroporation of the postsynaptic neuron, targeted by shadow-imaging with Alexa 594, in mice expressing GCaMP6 in excitatory neurons; imaging of the postsynaptic neuron, labelled by dsRed; injection of the modified rabies virus; imaging and tracing of the entire population, with presynaptic neurons marked by dsRed. (**b**) Time lapse of electroporation (Day 0) and dsRed expression (Day 1) of the postsynaptic neuron. Scale bar: 20 µm. (**c**) The postsynaptic neuron (*yellow*), its presynaptic ensemble (marked by dsRed, *red*) and the excitatory population (expressing GCaMP6, *green*). Lines indicate cortical layers. Montage of maximum intensity projections of structural z-stacks. Scale bar: 100 µm. (**d**) Example excitatory presynaptic neuron: expression of dsRed2 (*top*) provides a somatic outline matching the expression of GCaMP6 (*bottom*). (**e**) Same as in **d** for an example presynaptic inhibitory neuron, which does not express GCaMP6. (**f**) Responses to drifting gratings of 11 postsynaptic neurons. (**g**) Responses to drifting gratings of 12 example presynaptic neurons from L2/3 (*red*), L4 (*dark red*) or L5 (*brown*) synapsing onto the postsynaptic neuron highlighted in **f** (*orange*). (**h**) Average tuning across the 11 postsynaptic neurons in **f**, after alignment of their preferred stimulus direction to 0 deg. (**i**) Distribution of presynaptic preferred direction relative to postsynaptic preferred direction, pooled across all layers (n=11). Thin curves: fits to individual experiments; thick curve: average across experiments. (**j**) Same as **i**, for presynaptic ensembles within L2/3 (*top*, n = 11) and within L4 (*bottom*, n=9). **p<0.01 circular correlation, circular V-test, or one-way ANOVA.

As expected from many^4-7^ but not all^1,2^ previous observations, we found strong agreement between the visual preferences of the postsynaptic neuron and those of its excitatory presynaptic ensemble across layers (Figure 1f-j). In most experiments (12/13) the postsynaptic neuron responded to drifting gratings, and was typically selective for orientation and direction (Figure 1f). We could thus compare their selectivity with that of the visually responsive presynaptic neurons (Figure 1g). Presynaptic excitatory neurons were prevalently tuned to the orientation preferred by the postsynaptic neuron (Figure 1g-i): more than twice as many presynaptic neurons preferred the postsynaptic preferred orientation than the orthogonal orientation (p <0.01, one-way ANOVA and p <0.01 circular V-test). These effects did not result from local biases in neuronal preferences^30,31^, because they were absent in the remaining simultaneously recorded population (Supplementary Figure 4). This co-tuning was prevalent not only^4,7^ in L2/3, but also in L4 (Figure 1j; p <0.01 and p <0.01, one-way ANOVA; p<0.01 and p<0.05 circular V-test). Previous observations of different biases in these sources of input^1^ might be explained by unmeasured differences in inhibitory and excitatory neurons and by uncontrolled biases in local populations^30,31^. In addition to this tuning similarity, the excitatory presynaptic ensemble showed higher correlated variability than the other neurons (Supplementary Figure 5).

The excitatory and inhibitory ensembles providing input to a layer 2/3 pyramidal cell followed markedly different laminar and horizontal distributions (Figure 2a-l). Excitatory inputs were densest in L4 and spanned a large vertical and horizontal range, while inhibitory inputs dominated in L2/3 and clustered around the postsynaptic neuron (Figure 2a-c). These differences in distribution were consistent across presynaptic ensembles (Figure 2d-f). Hence, while measures of functional connectivity *in vitro* suggested global spatial balance between excitation and inhibition^27,28^, or even broader horizontal inhibition^29^, connectivity onto L2/3 neuron followed simple rules consistent with neuronal anatomy^23^. Vertically, L2/3 neurons received sparse excitation from L2/3, dense excitation from L4, and dense inhibition from L2/3 (Figure 2g-i). Horizontally, the broader distribution of excitatory relative to inhibitory inputs generated an inverse Mexican hat profile, with inhibition dominating proximal regions and excitation distal ones (Figure 2j-l, Supplementary Figure 6).

**Figure 2.**
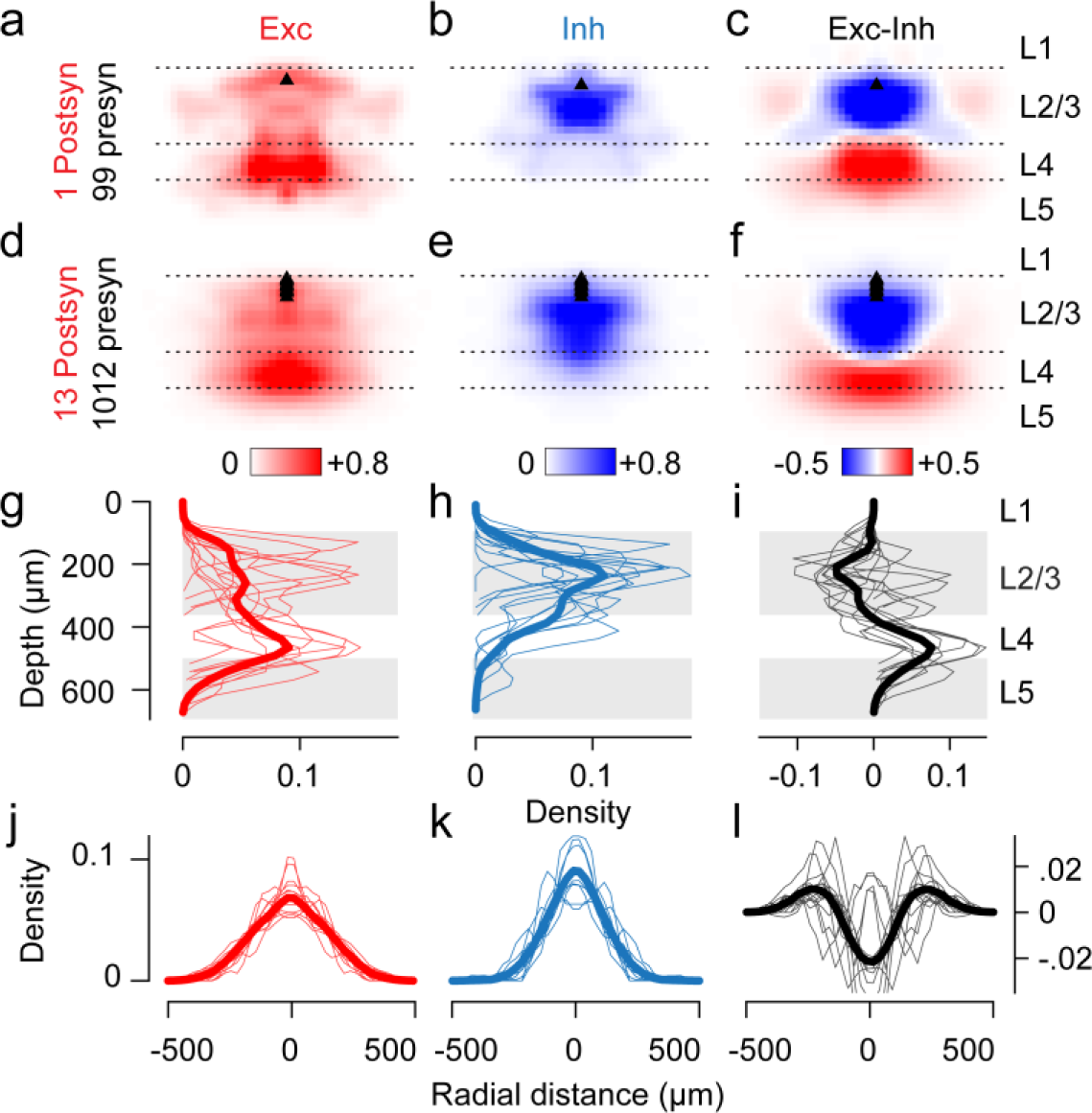
Anatomical distribution of excitatory and inhibitory presynaptic ensembles. (**a**) Horizontal and vertical density of excitatory presynaptic neurons around the postsynaptic neuron (*black triangle*), for the experiment shown in Figure 1c. Density was normalised to its maximum for display purposes. (**b**) Same as **a**, for inhibitory presynaptic neurons. (**c**) Difference between excitatory and inhibitory densities shown in **a** and **b**. (**d-f**) Same as **a-c** for data pooled across experiments (n=13). All postsynaptic neurons resided in L2/3 (*black triangles*). (**g-i**) Depth distributions for the probability densities in **a-c**, showing individual experiments (*thin lines*), and pooled data (*thick line*). Same vertical scale as in **a-f**. (**j-l**) Same, for the horizontal distributions. Same horizontal scale as in **a-f**.

We then asked if the differences in distribution between excitatory and inhibitory presynaptic neurons could support the selectivity of the postsynaptic cell (Figure 3a,b). To relate selectivity to anatomical distribution, we converted cortical positions to visual positions. We mapped the retinotopy of the imaged cortical volume using a sparse noise stimulus, and estimated the receptive field (RF) centre for every cortical location (Figure 3a). We could then assign RF centres to all neurons^9^, including unresponsive excitatory neurons and inhibitory neurons, which did not express GCaMP6 (Figure 3b). We used this retinotopic map to transform the location of a neuron from a point in cortical space (Figure 3a) to a point in visual space (Figure 3b).

**Figure 3.**
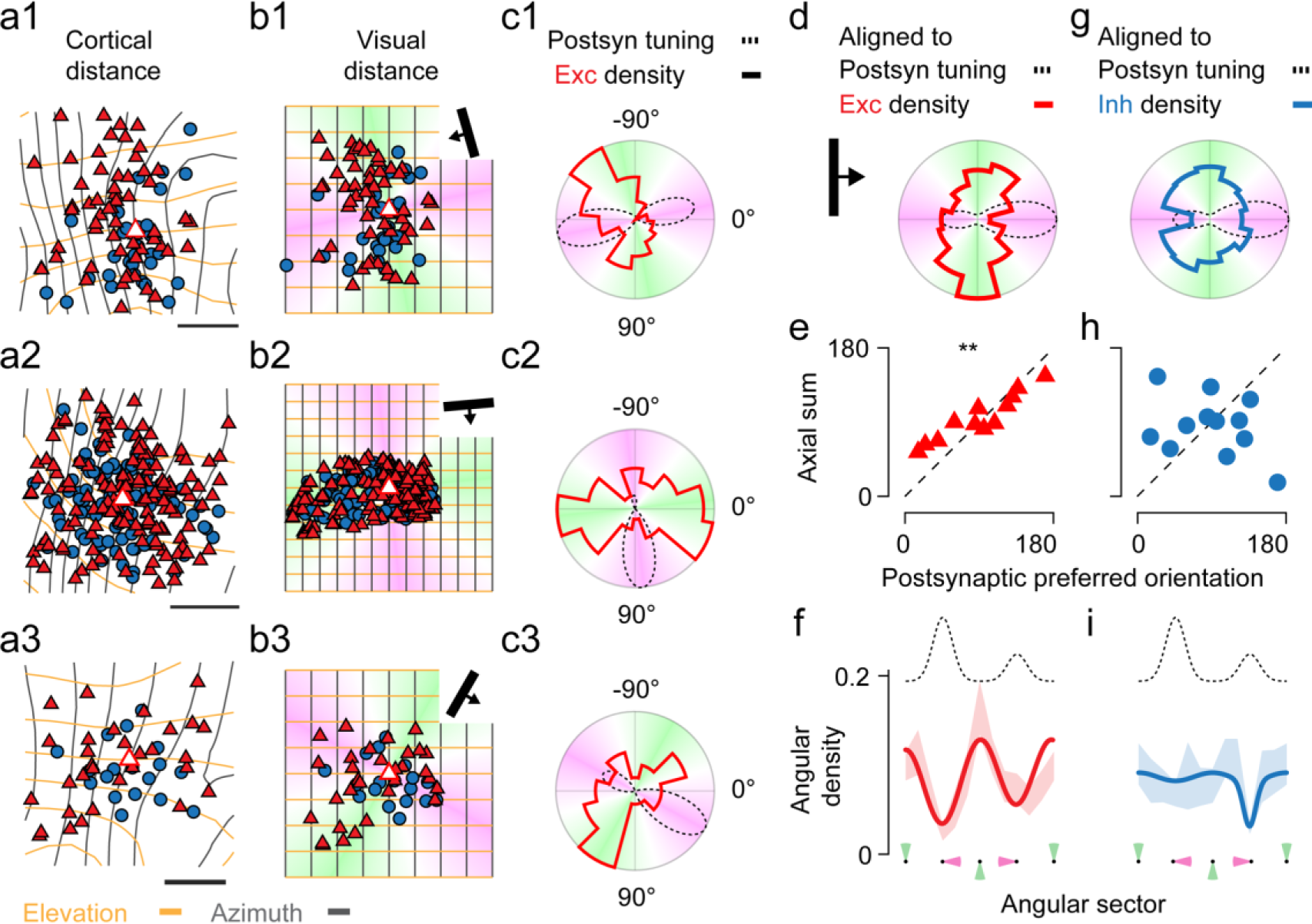
The presynaptic excitatory ensemble, but not the inhibitory one, is coaxially aligned with the orientation preference of the postsynaptic neuron. (**a**) Cortical distribution of excitatory presynaptic neurons (*red*) and inhibitory presynaptic neurons (*blue*) around an example postsynaptic neuron (*white fill*). Contours mark visual distance in azimuth (*grey*) and elevation (*orange*) from the RF of the postsynaptic neuron, in 5 deg steps. Scale bar: 200 µm. (**b**) The same cells as **a** replotted in visual space, so that contour lines describe a grid. Background colour denotes sectors of visual space around the RF of the postsynaptic neuron, which are orthogonal (*green*) or parallel (*magenta*) to the postsynaptic neuron preferred direction. Inset: preferred orientation and direction of the postsynaptic neuron (*bar* and *arrow*). (**c**) Polar representation of the angular density of excitatory presynaptic neurons (*red*), and of the postsynaptic neuron’s responses to drifting gratings (*black*). Background colour as in **b**. (**d**) Same as c, averaged across 12 experiments after alignment to the postsynaptic neuron’s preferred direction. (**e**) Axial vector sum of the distribution of the excitatory presynaptic ensembles, plotted against the preferred orientation of the postsynaptic neuron, for n = 12 experiments. Circular correlation 0.76, p = 0.01. (**f**) Angular density of the excitatory presynaptic neurons relative to the preferred direction of the postsynaptic neuron (*dotted tuning curve*). Insets label angular sectors orthogonal (*green*) or parallel (*magenta*) to the preferred direction. Data (*shaded area*, mean ± s.e., n= 12) were fit by the sum of a sinusoid (to capture orientation selectivity) and a Gaussian function (to capture direction selectivity). (**g-I**) Same as **d-f**, for inhibitory presynaptic ensembles. Circular correlation −0.06, p = 0.83.

The distribution in visual space of presynaptic excitatory neurons was elongated, along an axis that precisely matched the postsynaptic neuron’s preferred orientation (Figure 3b-f). The angular distribution of the excitatory presynaptic ensemble around the postsynaptic neuron showed marked axial asymmetries both in cortical and visual space (Figure 3b, c, Supplementary Figure 7). The orientation of the axial vector sum of these distributions was significantly correlated with the orientation preference of the postsynaptic neuron (Figure 3d, e; p<0.01, circular correlation, p<0.01 circular V-test). Indeed, excitatory inputs were most common in angular sectors that were parallel to the preferred orientation of the postsynaptic neuron (Figure 3c-d, f). Neither effect could be explained by the anisotropy of the retinotopic map^32^: presynaptic ensembles had significantly higher elongation and alignment than control surrogate ensembles, crafted to be isotropic in cortex and match the data in size and spatial scale (p <0.01 for both circular correlation and circular V-test; Supplementary Figure 7). This elongation and alignment of excitatory inputs were present throughout the postsynaptic RF, not only in the far periphery^5^, and would thus contribute to the cell’s selectivity to orientations presented within the RF.

Presynaptic inhibitory neurons, by contrast, were distributed more concentrically around the postsynaptic neuron and showed no alignment to its preferred orientation (Figure 3g, h). Excitatory presynaptic ensembles were significantly more elongated then inhibitory ones, both in cortical and visual space (Supplementary Figure 7, p<0.05 in cortex, p<0.01 in visual space, Wilcoxon signed-rank test). The asymmetry in the angular count of inhibitory neurons around the postsynaptic cell was less pronounced and was in many cases only as strong as expected by chance (Figure 3g, h, Supplementary Figure 7). Indeed, the vector sum of the axial distribution of inhibitory neurons around the postsynaptic cell did not significantly correlate with the orientation preference of the postsynaptic cell (Figure 3h; p>0.05, circular correlation and circular V-test). This contrast between elongated excitation and concentric inhibition could enhance the postsynaptic responses to the preferred orientation relative to other orientations. The presynaptic inhibitory neurons, however, were not distributed equally around the postsynaptic neuron; rather, they specifically avoided the angular sector upstream to the preferred direction of the postsynaptic neuron (Figure 3g, i). Excitatory neurons, instead, showed the opposite tendency (Figure 3d,f). This difference between excitation and inhibition suggested a possible mechanism for direction selectivity^14,19^. To examine how differences in angular density depended on the distribution of presynaptic neurons in different regions of the postsynaptic RF, we analysed the density of presynaptic neurons in visual space.

Excitatory neurons favoured visual locations upstream of the preferred direction, while inhibitory neurons favoured more downstream locations, centred closer to the postsynaptic neurons (Figure 4a-d). To combine the excitatory and inhibitory presynaptic densities measured in each experiment we rotated them, aligned them, and matched them in spatial scale to a common template in which the preferred direction of the postsynaptic neuron pointed rightwards (Figure 4a, b). The density of excitatory inputs in visual space was not only elongated along the orientation preference of the postsynaptic neuron, but also offset upstream to the neuron’s preferred direction of motion (Figure 4a, c). The density of inhibitory inputs, instead, concentrated around the postsynaptic neuron, with a smaller, yet consistent, offset downstream of the postsynaptic neuron’s preferred direction. (Figure 4b, c). This difference between excitation and inhibition was strongest amongst neurons located close to the postsynaptic cell, and gradually weakened in distal regions. (Figure 4c, d). To compare the inputs received by each postsynaptic neuron along its preferred direction, we calculated the density of excitatory and inhibitory neurons found in the angular sectors upstream and downstream of the postsynaptic RF (Figure 4c, e). Excitatory neurons were significantly more abundant upstream of the preferred direction, while inhibitory neurons dominated downstream (p<0.5, Wilcoxon signed-rank test). Indeed, the balance between the density of excitatory and inhibitory inputs invariably favoured visual sectors upstream of the RF (Figure 4c, f), particularly for the postsynaptic neurons with stronger direction selectivity (Figure 4e, f, p<0.01, Wilcoxon signed-rank test).

**Figure 4.**
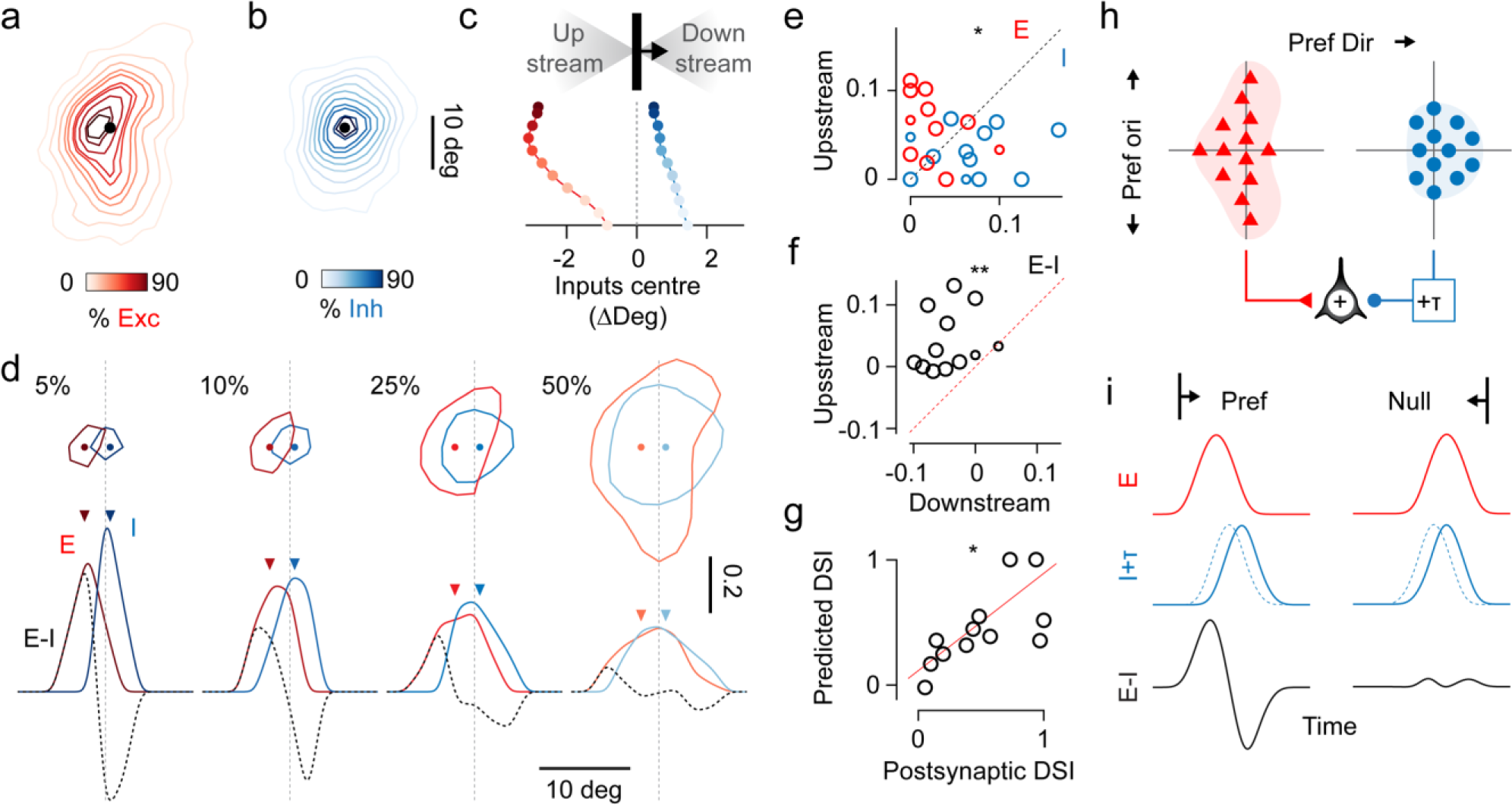
Spatially offset excitation and inhibition provide cortical mechanism for direction selectivity. (**a**) Density of excitatory presynaptic neurons as a function of RF distance from the postsynaptic neuron, rotated and averaged across experiments (*n=12*) according to the preferred orientation and direction of the postsynaptic neuron. Colormap indicates the percentage of total presynaptic neurons enclosed by each contour. (**b**) Same as in **a**, for inhibitory presynaptic neurons. (**c**) Azimuth centre of mass, relative to the RF of the postsynaptic neuron, of the excitatory (*red*) and inhibitory (*blue*) inputs enclosed by the contours in **a-b**, and lying in angular sectors of visual space upstream or downstream to the direction selectivity (*inset*). (**d**) Contour lines enclosing the central 5, 10, 25 and 50% of the excitatory and inhibitory presynaptic densities (*top*), and their sum projected on the axis of preferred direction (*bottom*). The azimuth centre of mass of each contour or distribution (dots and arrowheads) are calculated as in **c**. (**e**) Fraction of excitatory (*red*) or inhibitory (*blue*) inputs measured in angular sectors of visual space upstream or downstream to the direction selectivity of each postsynaptic neuron. Large circles represent postsynaptic neurons with DSI >0.1; small circles represent postsynaptic neurons with DSI ≤0.1. (**f**) Same as **e**, for the difference between the fraction of excitatory and inhibitory inputs measured in the upstream sector vs. the downstream sector. (**g**) The direction selectivity index predicted from the E-I asymmetries in **f** plotted against the direction selectivity index measured from the postsynaptic neuron (r =0.67, p <0.05). (**h**) Schematic of the organization of excitatory and inhibitory presynaptic ensembles: excitation coaxial with postsynaptic orientation preference, and asymmetric excitation and inhibition along the postsynaptic preferred direction. (**i**) If the inhibitory presynaptic neurons are activated after a delay (+τ), this model generates direction selective responses in the postsynaptic neuron

This spatial offset between excitation and inhibition could provide a circuit basis for direction selectivity (Figure 4g-i). Because inhibitory neurons in L2/3 must respond after excitatory neurons^12,33,34^, this offset can enforce direction selectivity^14,19,21,22^ (Figure 4h, i). When a moving stimulus approaches the postsynaptic RF in the preferred direction, excitation is recruited first and inhibition second, leading to a large postsynaptic response (Figure 4h, i). Conversely, in the opposite direction, inhibition is recruited first and its delay allows for the timely suppression of the postsynaptic response (Figure 4h, i). This mechanism would be sufficient to explain the direction selectivity of the postsynaptic neurons. Indeed, their direction selectivity could be predicted based on the spatial offsets of their excitatory and inhibitory presynaptic fields (Figure 4g, r = 0.66, p <0.05). While asymmetric excitation could, by itself, provide such offsets, the downstream displacement of inhibition suggests functionally specific inhibitory connections are instrumental to establish direction selectivity.

These results demonstrate that functionally specific differences in the spatial arrangement of excitatory and inhibitory connectivity can establish intracortical circuits for orientation selectivity and direction selectivity. The alignment of excitatory inputs could implement a mechanism for orientation selectivity similar to the alignment in visual space seen in thalamocortical circuits^15-19,35^, sharpened by concentric inhibition. The spatially offset arrangement of excitation and inhibition could provide a mechanism for direction selectivity, similar to the one observed for direction selective ganglion cells in the retina^21,22^. Layer 2/3, thus, replicates circuit motifs that appear at earlier stations of the visual system, and the reappearance of these motifs suggests that they might be canonical across brain regions.

## Acknowledgments

We thank Ed Callaway for sharing viruses via the Salk Institute Vector Core, Matteo Rizzi for advice on electroporation, Michael Krumin for support with microscopy, Charu Reddy for help with plasmid production, Massimo Scanziani and Pip Coen for suggestions on the manuscript. We also thank Kamill Balint, Stuart Trenholm, Botond Roska, and Marco Tripodi for advice and for genetic materials that we used in pilot studies. This work was supported by the Wellcome Trust (grants 099692/Z/12/Z to LFR; grants 205093 and 108726 to KDH and MC). MC holds the GlaxoSmithKline / Fight for Sight Chair in Visual Neuroscience.

## Author contributions

LFR, KDH and MC conceived the experiments. LFR refined the techniques, performed the experiments, and analysed the data. LFR, KDH and MC wrote the paper.

## Author information

Correspondence and requests should be addressed to.

## Methods

All experimental procedures were conducted in accordance with the UK Animals Scientific Procedures Act (1986). Experiments were performed at University College London under personal and project licenses released by the Home Office following appropriate ethics review.

### Transgenic mice breeding

Experiments were performed on fifteen 7-12 weeks old mice of both sexes, maintained on a 12 hours light/dark cycle. In these animals, GCaMP6 expression was achieved through different transgenic strategies. Six mice were Camk2a-tTA; EMX1-Cre; TIGRE-Ins-TRE-LSL-GCaMP6f (Ai93D) triple transgenic mice, expressing the calcium indicator GCaMP6f in all cortical excitatory neurons. Six mice were Camk2a-tTA; EMX1-Cre; TIGRE-Ins-TRE-LSL-GCaMP6s (Ai94D) triple transgenic mice, expressing the calcium indicator GCaMP6fs in all cortical excitatory neurons. Four mice were Camk2a-tTA; tetO-GCamp6s double transgenic mice, expressing GCaMP6s in all cortical excitatory neurons. Transgenic mice were bred from the following parental lines: Emx1-IRES-Cre (Stock #005628, The Jackson Laboratory, Ref. ^36^); CamK2a-tTA (Stock #007004, Ref. ^37^; Ai93 (Stock #024103, Ref. ^38^; Ai94 (Stock #024104, Ref. ^38^); TRE-GCamp6f (Stock #024742, Ref. ^39^). Differences in the GCaMP6 variants used did not affect the results of the study, because data in each experiment were compared against internal controls.

### Surgical procedures

The experiment entailed three surgical procedures (Figure 1a), each carried out under isoflurane anaesthesia (1-2% in Oxygen), while the body temperature was monitored and kept at 37-38 °C using a closed-loop heating pad, and the eyes were protected with ophthalmic gel (Viscotears Liquid Gel, Alcon Inc.). An analgesic (Rimadyl, 5 mg/kg) was administered subcutaneously on the day of the surgical procedure and on subsequent days, as needed. Whenever the procedure exposed the brain, Dexamethasone (0.5 mg/kg, IM) was administered intramuscularly 30 min prior to the procedure to prevent brain oedema. The exposed brain was perfused with artificial cerebrospinal fluid at all times (150 mM NaCl, 2.5 mM KCl, 10 mM HEPES, 2 mM CaCl_2_, 1 mM MgCl_2_; pH 7.3 adjusted with NaOH, 300 mOsm).

The first surgery consisted of the implant of a stainless steel head-plate over the right hemisphere of the cranium. The head was shaved and disinfected; the cranium was exposed and covered with biocompatible cyanoacrylate glue (Vetbond). A stainless-steel head plate with a 10 mm circular opening was secured over the skull using dental cement (Super-Bond C&B, 10 Sun Medical Co. Ltd., Japan). The exposed bone inside the chamber was covered by a thin layer of dental cement and sealed off with silicone elastomer (KwicKast). The animal was then allowed to recover for at least 4 days before further experimental procedures.

The second surgery was aimed at targeted neuron electroporation. A 1.5-2 mm wide square craniotomy was opened over the visual cortex (centred at −3.3mm AP, 2.8 ML from bregma). The animal was then transferred to the two-photon microscope setup for single neuron electroporation. Finally, the craniotomy was sealed with a glass cranial window, attached to the skull using cyanoacrylate glue and dental cement. The window was assembled from a circular cover glass (3 mm diameter, 100 µm thickness) glued to a smaller custom made glass square insert (1.5-2 mm wide, 300 µm thickness) with index-matched UV curing adhesive (Norland #61).

After electroporation and imaging of the postsynaptic cell, we performed a third surgery to inject a rabies virus (RV) for single neuron initiated monosynaptic tracing^40-43^. The cranial glass window was removed^44^ and a durotomy performed using a recurved insulin needle. Then, 100-200 nL of EnvA-dG-dsRed2-RV (10^8^-10^9^ pfu/mL) was injected using a pneumatic injector (Nanoject, Drummond Scientific Company) through a 30-50 µm borosilicate capillary. The injection was targeted ~100-200 µm from the electroporated neuron, with the help of reference images of the brain vasculature. Finally, a permanent glass window was implanted as described before.

In control experiments, we injected two mice with a combination of diluted AAV2.1-CaMK2a-Cre (~10^7^ GC/mL) and concentrated AAV2.1-flex-Syn-tdTomato (~10^12^ GC/mL) to express the red fluorescent protein tdTomato in a sparse ensemble of excitatory neurons.

### Targeted single cell electroporation with survival control

Targeted, single-cell electroporation^45,46^ was performed under a Sutter-MOM two-photon microscope, equipped with a low magnification (16X) high NA (0.8) water immersion objective lens (Nikon), and a femtosecond pulsed laser (Chameleon Ultra II, Coherent) tuned at 820nm. The microscope was controlled using ScanImage v3.8 (Ref. ^47^). We used an epifluorescence imaging module and camera to take reference pictures of the vasculature and guide the electroporation to a brain region devoid of blood vessels.

Borosilicate pipettes (resistance 10-14 MΩ) were crafted with a vertical pipette puller (Narishige) and filled with intracellular solution (133 mM KMeSO_4_, 7 mM KCl, 10 mM HEPES, 2 mM Mg-ATP, 2 mM Na_2_-ATP, 0.5 mM Na_2_-GTP, 0.05 mM EGTA; pH 7.2, adjusted with KOH, 280–290 mOsm; filtered using a 0.45 mm syringe filter). The intracellular solution also contained 50 µM of the red fluorescent dye AlexaFluo 594 and 3 plasmids with the following concentrations^42^: 100µg/µL pCAG-dsRed2 (Addgene #15777), 200 µg/µL pCMV-oG^48^ (Addgene 74288) (or pCMV-G in a subset of experiments, Addgene 15785) and 100 µg/µL pCAG-TVA800(Addgene 15788).

The pipette was manoeuvred with a micromanipulator (Junior 4 axis, Luigs&Neumann) and pushed through the dura, while applying positive pressure (~150 mBar) and monitoring the pipette resistance with an electroporator for *in vivo* transfections (Axoporator 800A, Molecular Devices). The pressure was then reduced to 30-50 mBar: the diffusion of the red dye in the extracellular space counterstained neuronal somas, while GCaMP6 allowed targeting of excitatory cells (Figure 1b). The pipette tip was pushed onto a neuronal soma, and, upon a ~30% increase in resistance, neurons were electroporated with a single pulse train at −11 V, 100 Hz, 0.5 ms pulse width, 1 s duration, and the pipette swiftly retracted.

Three criteria were used as signature of successful electroporation and recovery of a neuron from the electroporation shock (Supplementary Figure 1). First: immediate filling of the soma with AlexaFluo594 (Supplementary Figure 1d-e). Second: retraction of the pipette without pulling of the neuronal membrane (Supplementary Figure 1d). Third: sustained somatic GCaMP6 fluorescence at 820 nm relative to the surrounding neuropil (Supplementary Figure 1g-i). Electroporated neurons were monitored for the 3-10 min post-electroporation. The somatic fluorescence was normalized to the fluorescence of the surrounding neuropil and neurons. This procedure gauged both the intracellular dynamics of GCaMP6 and corrected for varying imaging quality during the time lapse. With these criteria, we could predict which electroporation attempts were successful with an 80% success rate.

GCaMP6 fluorescence at 820 nm was the best predictor of cell integrity and neuronal recovery after electroporation. At this wavelength, the calcium-bound and calcium-free isoforms of GCaMP6 are approximately equally fluorescent, as 820nm represent an isosbestic point of the two-photon cross section of the GCaMP6 isomers (unpublished 2P action cross section spectra from Harris Lab website, Janelia Farm). Hence, GCaMP6 fluorescence at 820 nm reports the intracellular concentration of the sensor. A healthy neuron should maintain its intracellular concentration of calcium indicator, and its somatic fluorescence should remain constant; conversely, in a neuron with irreversible membrane damage, the indicator should gradually diffuse outside of the cell, and the neuron should darken and disappear against the surrounding neuropil, where GCaMP6 concentration is assumed instead constant (Supplementary Figure 1i).

We performed up to five electroporation attempts per mouse, at sites spaced >500 µm apart. Transgene expression of electroporated cells was assessed 1-3 days after electroporation (Supplementary Figure 1j).

### Targeted photoablation of supernumerary electroporated neurons

In experiments where multiple neurons survived the electroporation, the functional characterisation of their visual responses allowed the selection for monosynaptic tracing of the neuron with the most desirable visual preference, namely significant orientation selectivity and retinotopic location centred in one of the stimulation screens. Supernumerary neurons were eliminated with targeted photoablation (Supplementary Figure 2). Photoablation was performed under the same setup used for electroporation. The galvanometric mirrors where centred on the target neuron’s soma, and pulses of high intensity (>100 mW) illumination were delivered for 10-20 s. Neurons were imaged between the pulses to titrate the extent of the photodamage (Supplementary Figure 2a-b). Photoablation pulses were delivered at 820 nm to restrict photodamage to the target neurons. At this wavelength, the two-photon cross-section of dsRed2, expressed only by the electroporated cells, is much higher than the two-photon cross section of GCaMP6, expressed by all neurons. Since photobleaching of the fluorophore is a major determinant of photodamage^49,50^, electroporated neurons were more sensitive to our photoablation protocol, and surrounding cells were rarely damaged. Successful photoablation was ascertained the following day (Supplementary Figure 2c). The injection of the rabies virus for monosynaptic tracing was withheld until the confirmed disappearance of supernumerary electroporated neurons.

### Two-photon imaging of neuronal responses

Recordings of neuronal activity were performed with a standard resonant-scanning two-photon microscope (B-Scope, Thorlabs), equipped with a Nikon 16x, 0.8 NA objective mounted on a piezoelectric z-drive (PIFOC P-725.4CA, Physik Instrumente, range 400 µm) for volumetric multi-plane imaging. The microscope was controlled using ScanImage v4.2 (Ref. ^47^). Excitation light was provided by a femtosecond laser (Chameleon Ultra II, Coherent), tuned between 920-1000 nm depending on imaging depth. Laser power was depth-adjusted between 30-300 mW, and synchronized with piezo position using an electro-optical modulator (M350-80LA, Conoptics Inc.). Sample fluorescence was collected in a green and a red channel: 525/50 nm for the GCaMP6, and 605/70 for the dsRed2. The imaging objective and the piezoelectric element were light shielded using a custom-made metal cone, a tube, and black cloth to prevent contamination of the fluorescent signal by the visual stimulation light.

Fields of view (FOV) were imaged with a resolution of 512*512 pixels at 30 Hz. FOV typically spanned 150-200 µm for the initial imaging of the postsynaptic neuron, and between 500-850 µm for the imaging of presynaptic ensembles. For volumetric imaging of GCaMP6s, the objective was scanned across 10 planes, separated by 15-20 µm in depth, resulting in an effective sampling rate of 3 Hz. For volumetric imaging of GCaMP6f, only 5 planes were used instead, with an effective sampling rate of 6 Hz. Serial volumetric imaging of the cortical volume around the postsynaptic cell was achieved by repeating volumetric acquisitions from the cortical surface to L5, using the piezo or the microscope stage motors to lower the objective between acquisitions.

During the imaging sessions, mice were head fixed on an airflow-suspended spherical treadmill. Each postsynaptic neuron and presynaptic ensemble were imaged daily, from the day following electroporation up to two weeks post rabies virus injection. These sparsely labelled neurons were used as a reference to target the same cortical volume during this longitudinal imaging.

To acquire structural z-stacks, the piezoelectric z-drive was used to move the objective in steps of 1 µm and image 10-500 repetitions of the same cortical volume. To extend the range of the piezoelectric z-drive, the procedure was repeated after moving the objective 400 μm down using the microscope motors. Z-stacks were used to quantify the total number of presynaptic neurons traced across days, align functional recordings to the reconstructions of the postsynaptic neuron, and identify borders between cortical layers in each experiment based on changes in neuronal morphology and neuronal density (Figure 1b).

### Visual stimulation

Visual stimuli were generated in Matlab (MathWorks) using the Psychophysics Toolbox^51^ and displayed on 3 gamma-corrected LCD monitors (refresh rate 60Hz) arranged at 90 degrees to each other. The LCD screens were covered with Fresnel lenses to correct for viewing angle inhomogeneity of the LCD luminance^52^. The mouse was positioned at the centre of the C-shaped monitor arrangement at 20 cm from all three monitors, so that the monitors spanned ±135 degrees of horizontal and ±35 degrees of the vertical visual field.

Sparse, spatial white noise stimuli were used to map the retinotopy of the imaged area and estimate the RF of neurons. Patterns of sparse black and white squares (4.5-6 degrees of visual field) on a grey background were presented at 5 Hz, typically in 10 min sequences repeated 3 times for each FOV. At any point in time, each square had a 2-5% probability of being non-grey, independent of the other squares.

To measure direction and orientation tuning we presented gratings drifting in twelve different directions, centred on the RF of the postsynaptic neuron. Gratings were presented in a circular window of 30-60 degrees, depending on the size of the imaged FOV, at 100% contrast on a grey uniform background, with a spatial frequency of 0.05 cycles per degree, and a temporal frequency of 2 Hz. At least ten repeats of each stimulus were presented for each FOV. Stimuli were 1-2 s in duration, and interleaved by 3-4 s of grey background.

### Processing of two-photon data

Two-photon data were pre-processed using Suite2p (Ref. ^53^). The pipeline included image registration, segmentation of active region of interest (ROIs), and estimation of the neuropil signal contaminating each ROI. The final selection of ROIs was curated manually to include only neuronal somas, and discard spurious or noisy ROIs. Active presynaptic neurons expressing dsRed2 were identified by inspecting the average dsRed2 image for each acquisition.

The neuropil signal was weighted by a correction factor α, determined separately for each ROI^54^, before being subtracted. The correction factor was estimated from the linear relationship between the lowest somatic fluorescence compatible with any value of fluorescence in the neuropil. For each neuron *i* the neuropil signal *N_i_(t)* was binned into 20 intervals; for each interval, the 5th percentile of the matching time-points of raw somatic fluorescence *F_i_(t)* was measured as an estimate of baseline fluorescence. α_i_ was then computed by linear regression between the median of each neuropil interval and estimates of baseline fluorescence, to accurately fit the lower envelope of the scatterplot of neuropil versus somatic fluorescence. The corrected fluorescence was computed as *F_i_ (t) – α_i_N_i_ (t)* and z-scored for further analysis.

Dense volumetric imaging often resulted in the same neurons being imaged multiple times in different imaging planes: duplicates of the same neuron were identified as ROIs whose centre of mass was closer than 5 µm in lateral distance and closer than 20 µm in depth, and had a signal higher than 0.5. Amongst duplicates, the ROI with the highest SNR was chosen for further analysis.

### Classification of excitatory and inhibitory neurons

Serial volumetric imaging time-series were averaged to obtain high signal to noise ratio z-stacks, with a green GCaMP6 channel, highlighting GCaMP6 expressing neurons, and a red dsRed2 channel, highlighting presynaptic neurons. Then, a custom algorithm was used to identify and classify presynaptic neurons either as excitatory or inhibitory. First, an iterative thresholding algorithm was used to segment somatic masks of presynaptic neurons from the red channel; somatic masks were inspected and curated manually, and spurious or out-of-focus neurons discarded (Supplementary Figure 3a, b). Then, the phase correlation (i.e. whitened cross-correlation) between the somatic mask of each neuron and its GCaMP6 fluorescence image was used to classify neurons as GCaMP6 positive or GCaMP6 negative units (Supplementary Figure 3a-d). The amplitude of the central correlation peak and the standard deviation of the correlation values inside a 5 µm annulus around the peak were used to fit a bilinear classification boundary separating putative excitatory GCaMP6 expressing neurons from putative inhibitory GCaMP6 negative neurons (Supplementary Figure 3c). This classification was computed independently for each experiment; the pooled results, with the average classification boundary, are presented in Supplementary Figure 3. The algorithm was tested in control experiments were a red protein was sparsely expressed only in excitatory neurons: in these datasets, the rate of false negatives (i.e. neurons wrongly classified as inhibitory) was below 5% (Supplementary Figure 3e-f).

The cortical position of presynaptic neurons was used to estimate their radial and layer densities around the postsynaptic neuron. Cortical probability densities were calculated in 625 µm^2^ bins and smoothed with a 50 µm wide Gaussian kernel. As described below, retinotopic maps were then used to transform the location of a neuron from a point in cortical space to a point in visual space. Retinotopic probability densities were calculated in 4 deg^2^ bins and smoothed with a 4 deg wide Gaussian kernel.

### Analysis of neuronal responses

All neurons identified by Suite2P were considered visually responsive and included for further analysis. Stimulus triggered responses were computed as the difference between the z-scored fluorescent trace and the 20^th^ percentile of the baseline activity 1s prior to stimulus presentation. Stimulus triggered average responses and standard errors were obtained across responses to the same type of stimuli.

Responses to drifting gratings were quantified as the integral of the fluorescent response in a 3 s window following the onset of stimulation. The preferred orientation and direction of motion were defined according to the grating eliciting the maximal response. Only for the postsynaptic neuron, the response window was optimised and the preferred direction and orientation were computed by fitting a double-Gaussian tuning curve to the data. The direction selectivity index of postsynaptic neurons was computed as:

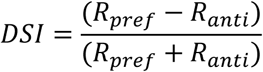

where *R*_*pref*_ is the response to the preferred direction, and *R*_*anti*_ is the response to the anti-preferred direction.

Neuronal RF centres were estimated from population retinotopic maps. For this analysis, imaging time series were denoised and compressed by SVD. First, a global receptive field was fit to the whole imaged volume: a single fluorescence time course was computed by averaging fluorescence from pixels within each frame, and concatenating in time frames from different planes. Before concatenation, the fluorescent intensity from different planes was equalised to the median signal throughout the recording. A spatiotemporal RF was calculated by reverse correlation between the signal and the absolute changes in luminance of the sparse noise stimulus, and then averaged in time to obtain a spatial RF. A global RF field centre was calculated as the centre of mass of the 10% most responsive pixels in the spatial RF. Second, the imaged volume was binned in quadrants using a 10 by 10 grid, and a new RF centre fit to individual quadrants; these fits were constrained to be within 60 deg in azimuth and 35 deg in elevation from the global preferred retinotopic location. RF centres from each quadrant were combined into a coarse retinotopic map, which was finally interpolated to the full pixel size of the FOV and smoothed with a 50μm Gaussian. The retinotopic map was used to transform between cortical coordinates to visual space coordinates and assign RF centres to each neuron based on their position in the FOV. For each dataset, the best recording was chosen as a reference retinotopy, and data from other recordings aligned to it. Retinotopic maps were also used to confirm the targeting of the experiment to V1, and discard from further analysis cortical territories in higher visual areas.

Pairwise correlations were estimated using Pearson’s correlation coefficient (Supplementary Figure 5). Simultaneously recorded neuronal responses from each imaged volume were aligned to a common time reference by interpolation. Total correlations were computed from neuronal responses over the duration of the whole recording, including periods of stimulation and spontaneous activity (Supplementary Figure 5a, b). Signal correlations were calculated from the average responses across trials of drifting gratings. Noise correlations were computed by subtracting the average response across trials from the responses in each trial and then calculating the correlation coefficient of the mean-subtracted responses over the whole recording. Total and noise correlations could be computed only for simultaneously recorded pairs, while signal correlations were computed for every recorded pair across sessions. The distribution of pairwise correlations between pairs of presynaptic neurons was compared to the distribution of pairwise correlations between pairs of other neurons using a one-tailed Kolmogorov-Smirnov test (Supplementary Figure 5c, d). We performed additional statistical analysis to control for biases between the presynaptic ensemble and the surrounding population. To control for the effects of proximity, we sorted neuronal pairs according to their lateral distance, and computed the median pairwise correlation in 50 µm bins (Supplementary Figure 5d, e). Moreover, given the difference in sample size between the two distributions, the significance of the Kolmogorov-Smirnov test result was compared against a null distribution obtained from 10^4^ size matched surrogate distributions, drawn from other simultaneously recorded neurons (Supplementary Figure 5f-h). Finally, average values of pairwise correlation were obtained for each presynaptic ensemble and compared to the average pairwise correlation across the population using the Wilcoxon sign-rank test. The same analysis was also repeated after grouping neurons according to the parent cortical layer and considering each layer ensemble as a separate sample (Supplementary Figure 5f-h).

### Orientation tuning and retinotopic alignment of presynaptic ensembles

Distributions of preferred direction and orientation were computed in 30 deg bins from all responsive neurons and centred to represent relative deviations from the postsynaptic selectivity. Multiple statistical tests were used to ascertain if presynaptic ensembles were significantly tuned to the selectivity of the postsynaptic neurons. First, variations in neuronal counts at different relative directions and orientations across datasets were tested with one-way ANOVA. Then, the preferred direction and orientation of presynaptic distributions, and their surrogate controls, was computed as the vector sum of each distribution. Then, two measures circular alignment were used to assess if the tuning of the presynaptic ensemble matched the tuning of the postsynaptic neuron. Circular correlation was used to measure how well the tuning of the presynaptic ensembles correlated with the tuning of the postsynaptic neurons. The circular V-statistic instead was used to assess if the distance between the selectivity of the postsynaptic neurons and their parent presynaptic ensemble was uniformly distributed, or rather formed a unimodal circular distribution with mean direction equal to 0. Finally, to control for local biases in the population, the data from presynaptic ensembles were compared against 10^4^ surrogate distributions matched in size, comprised of other simultaneously recorded neurons. The circular correlation coefficient and the V-statistic of presynaptic ensembles were tested against null distributions computed from the surrogate ensembles to obtain an empirical p-value. This analysis was performed either by including all presynaptic neurons, or by splitting them by layer based on cortical depths. Surrogate distributions were generated accordingly. Angular distributions in visual space were fit as

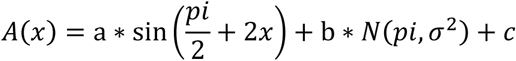

where the density *A* of presynaptic neurons in each angular bin *x* around the postsynaptic neuron was modelled as the sum of a sinusoidal function aligned to the postsynaptic preferred orientation and a Gaussian function aligned to the postsynaptic anti-preferred direction.

The circular variance of the angular and axial distribution of excitatory and inhibitory presynaptic neurons around the postsynaptic neurons was calculated to measure cortical or retinotopic elongation (Supplementary Figure 7). Angular and axial distributions were computed in 30 deg bins, both in cortical and retinotopic space. These distributions were compared against 10^4^ surrogate angular and axial symmetric distributions, matched in size to the data. The circular variance of these surrogate distributions was used to build a null distribution and obtain a statistic for the significance of the axial and angular anisotropy of presynaptic ensembles. The smallest of the two was used as a p-value for the elongation of the data against the null hypotheses of symmetry (Supplementary Figure 7a-e), and the log_10_(p-value) used as a measure of anisotropy.

As a further control, presynaptic ensembles were modelled as 2D isotropic Gaussians in cortex: these fits were used to craft 10^4^ surrogate retinotopic ensembles, matched in size and spatial scale to excitatory or inhibitory ensembles. These distributions were isotropic in cortical space by design, and once transformed into visual space, reflected and controlled for biases due to local inhomogeneity or varying magnification factor in the retinotopic map. (Supplementary Figure 7f).

Using the latter surrogate distributions as controls, the alignment between the retinotopic elongation of presynaptic neurons and the selectivity of the postsynaptic cell was tested with circular correlation and circular V-test, with analyses equivalent to the ones described for the presynaptic distributions of direction and orientation preference.

To average the excitatory and inhibitory presynaptic densities measured in each experiment, we rotated them, aligned them, and scaled them to a common template in which the preferred direction of the postsynaptic neuron pointed rightwards. Rotation was designed to align the preferred direction of the postsynaptic cell to 0 deg in visual space. Scaling was tailored to match in size symmetric Gaussian fits to the density of the inhibitory ensembles. Using the same fits, alignment was designed to remove jitter in elevation.

To estimate the direction selectivity of the postsynaptic neuron from the distribution of excitatory and inhibitory presynaptic ensembles, we used the following expression:

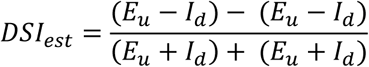

where *E*_*u*_ and *I*_*u*_ are the densities of excitatory and inhibitory presynaptic neurons in the upstream angular sector, and *E*_*d*_ and *I*_*d*_ are the same, for the downstream angular sector.

Circular statistics were computed using the CircStat toolbox^55^ in Matlab (MathWorks).

---

**Supplementary Figure 1.**
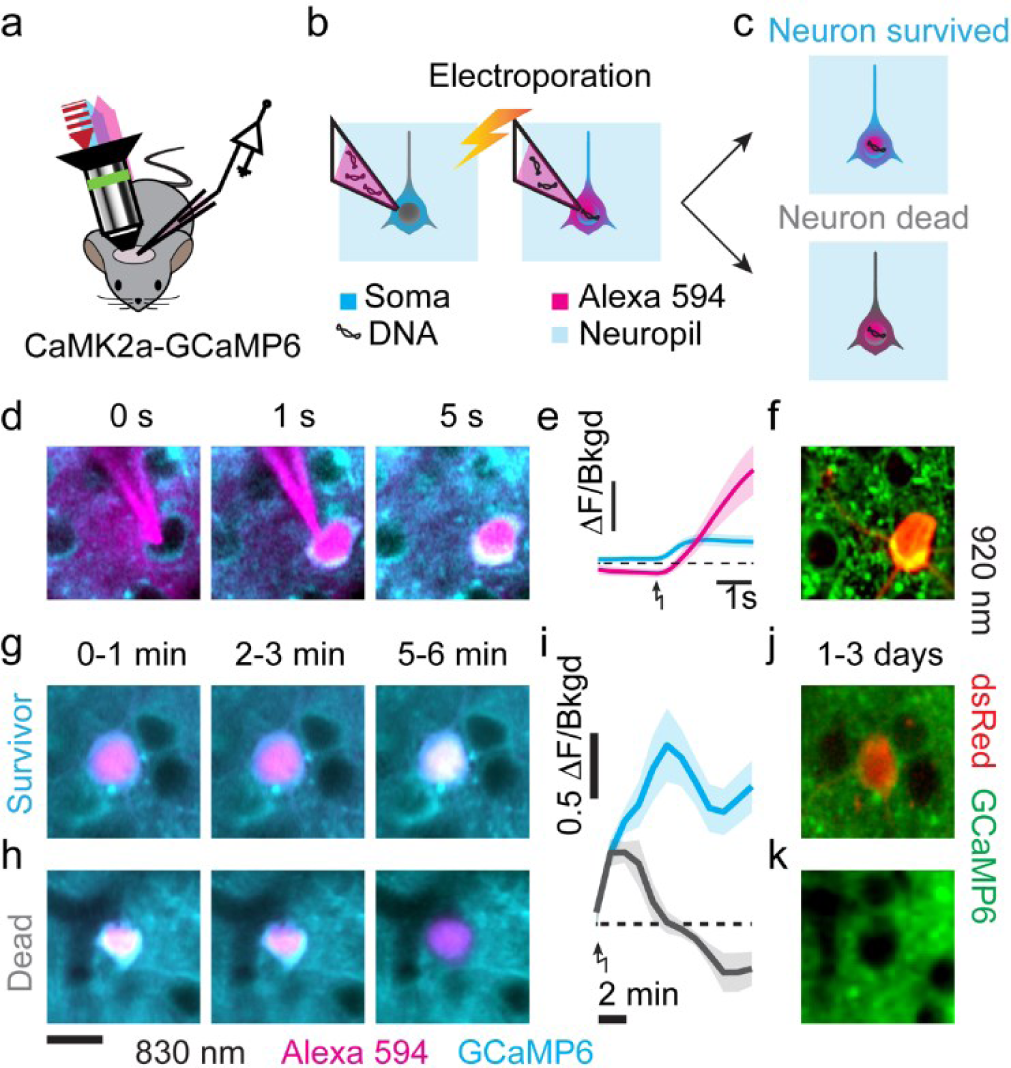
Targeted single neuron electroporation with survival control *in vivo*. (**a**) Electroporation was performed on a transgenic mouse expressing GCaMP6 in cortical excitatory neurons. A pipette, filled with Alexa 594, is targeted to a craniotomy; an 820 nm laser (*red*) excites Alexa 594 fluorescence (*magenta*) and calcium insensitive GCaMP6 fluorescence (*cyan*). (**b**) Upon electroporation, DNA plasmids and Alexa594 are transferred into a neuron expressing GCaMP6. (**c**) The comparison between the intensity of GCaMP6 fluorescence in the soma and the surrounding neuropil is predictive of neuron survival. Neurons that survive maintain GCaMP6 fluorescence (*top*), while irreversibly damaged neurons disappear against the neuropil background (*bottom*). (**d**) Time-lapse of the electroporation in layer 2/3 of mouse V1, using Alexa 594 negative contrast and calcium insensitive GCaMP6 fluorescence imaging: approach (*left panel*), electroporation (*middle panel*), pipette withdrawal (*right panel*). (**e**) Time-course of somatic Alexa 594 (*magenta*) and GCaMP6 (*cyan*) fluorescence relative to neuropil background (*dashed line represents unity*) during electroporation (*arrow*, n = 10). (**f**) The neuron in **e**, imaged at 920 nm, expressing the electroporated genes for dsRed (*red*) and maintaining healthy GCaMP6 expression (*green*) 1 day after the procedure. (**g**) Same as **d**, for a 5 min long time-lapse of a neuron that survived the procedure. Images are 30 s long averages acquired between 0-1 min (*left*), 2-3 min (*centre*) and 5-6 min (*right*) after electroporation. (**h**) Same as **g**, for a neuron that did not recover from the electroporation. (**i**) Same as **e,** for GCaMP6 somatic fluorescence of neurons that did (*cyan, n=18*) or did not (*grey, n=10*) survive the procedure. (**j**) Same as **f**, for the neuron shown in **g**. (**k**) Same as **f**, for the neuron shown in **h**. Scale bar 30 µm, same for all fluorescence images.

**Supplementary Figure 2.**
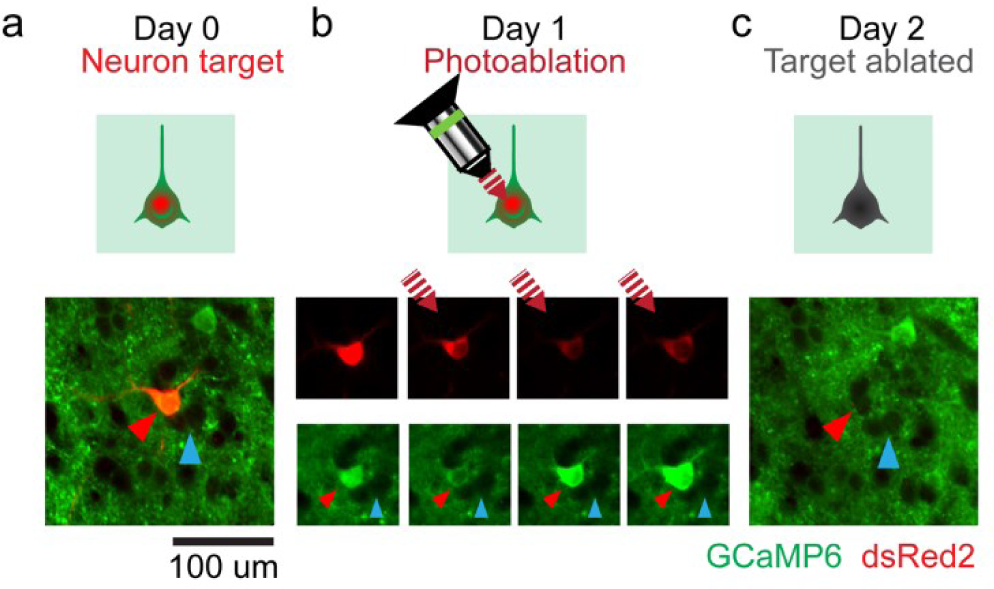
Photo-ablation of supernumerary postsynaptic neurons before rabies injection. (**a**) The target neuron (*top and bottom*) expresses both GCaMP6 (*green*) and dsRed (*red, red arrow*), while surrounding neurons (*cyan arrow*) only express GCaMP6. (**b**) The photo-ablation procedure: scanless two-photon illumination at 820 nm (*red dashed line*) is focused on the neuron, with intensity > 100 mW, for 10-20 seconds (*top*). Time lapse imaging at 920 nm of the target neuron between the photo-ablation pulses (*bottom*), showing increasing signs of localized cellular photo-damage: elevated intracellular calcium, localised photo-bleaching and cell swelling (*red triangle*). Neighbouring cells, not expressing the dsRed, resist the photo-damage (*cyan arrow*). (**c**) The successful photo-ablation of the target neuron (*top and bottom*) resulted in its complete disappearance on the day following the procedure (*red arrow*), while surrounding cells remained unaffected (*cyan arrow*).

**Supplementary Figure 3.**
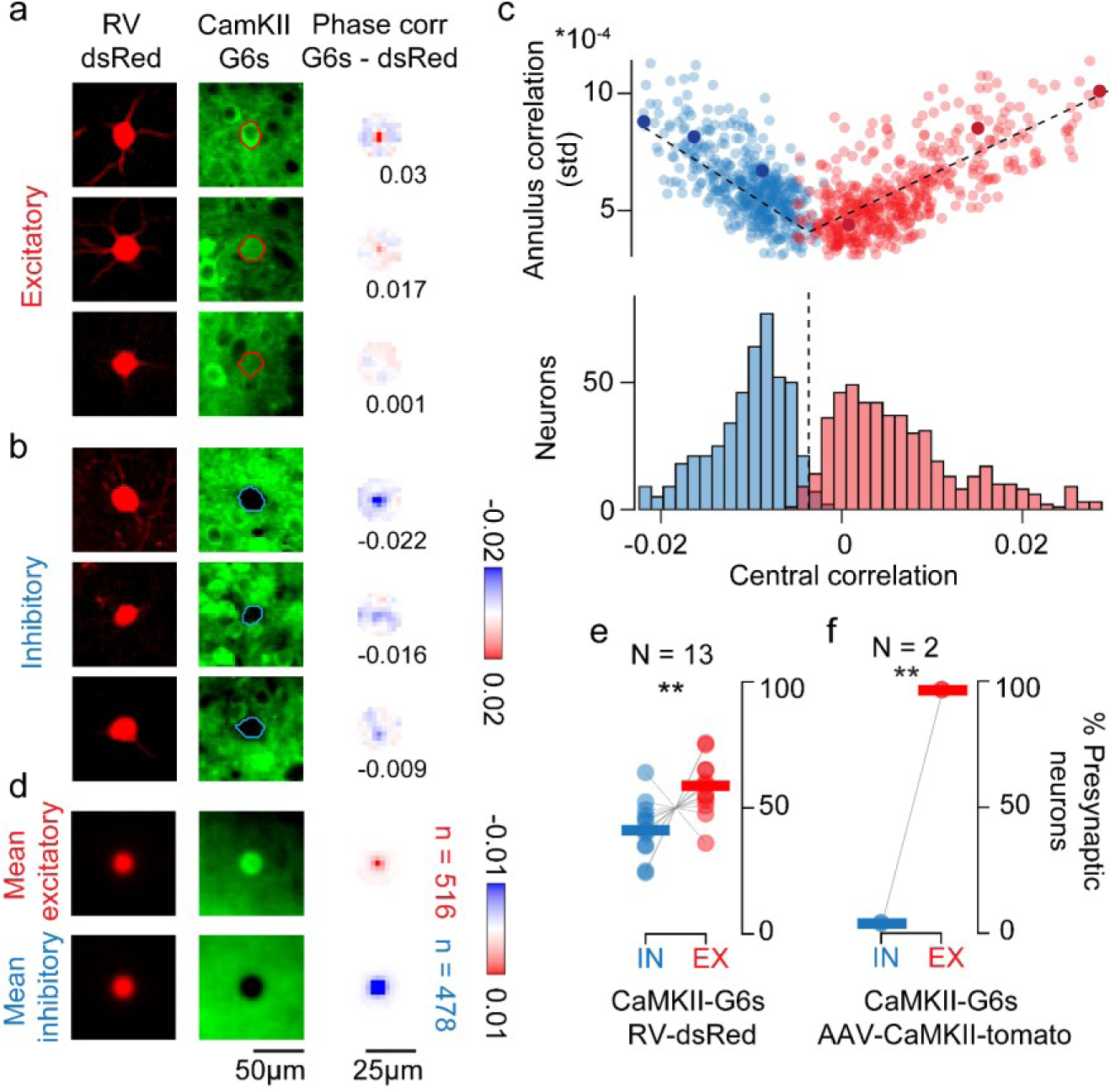
Classification of excitatory and inhibitory presynaptic neurons. (**a**) Three example excitatory presynaptic neurons: expression of dsRed (*left*) provides a somatic outline matching the expression of GCaMP6 (*middle*). The somatic mask and the GCaMP6 signal were used to compute a map of phase correlation in the 25 μm annulus around the somatic centre (*right*). The positive amplitude of the central peak of the phase correlation was used as a proxy for the intensity of GCaMP expression in the soma compared to the surrounding neuropil. (**b**) Same as **a** for three example inhibitory neurons. The negative amplitude of the central phase correlation indicated lack of GCaMP6 expression in the soma. (**c**) For each presynaptic neuron, the central peak of phase correlation was plotted against the standard deviation of the phase correlation within the 25 μm annulus around the soma (*top*). The cluster of excitatory neurons (*red*) and inhibitory neurons (*blue*) were split using a classification boundary identified with a bilinear fit to the data (*black line*). The plot displays a summary of neuronal classification across experiments, with the average classification boundary (*bottom*). (**d**) Average expression of dsRed (*left*), GCaMP6 (*middle*) and map of phase correlation (*righ*t) for presynaptic neurons classified as excitatory neurons (*top,* n=516) or inhibitory neurons (*bottom*). (**e**) Percentage of excitatory (*red*) and inhibitory (*blue,* n = 478) neurons in each presynaptic ensemble (*grey lines indicate paired data*), with median across experiments (*thick lines*). (**f**) Same as **e**, for experiments where a red marker, tdTomato, was expressed only in excitatory neurons. The rate of false negatives classified as inhibitory neurons was below 5%.

**Supplementary Figure 4.**
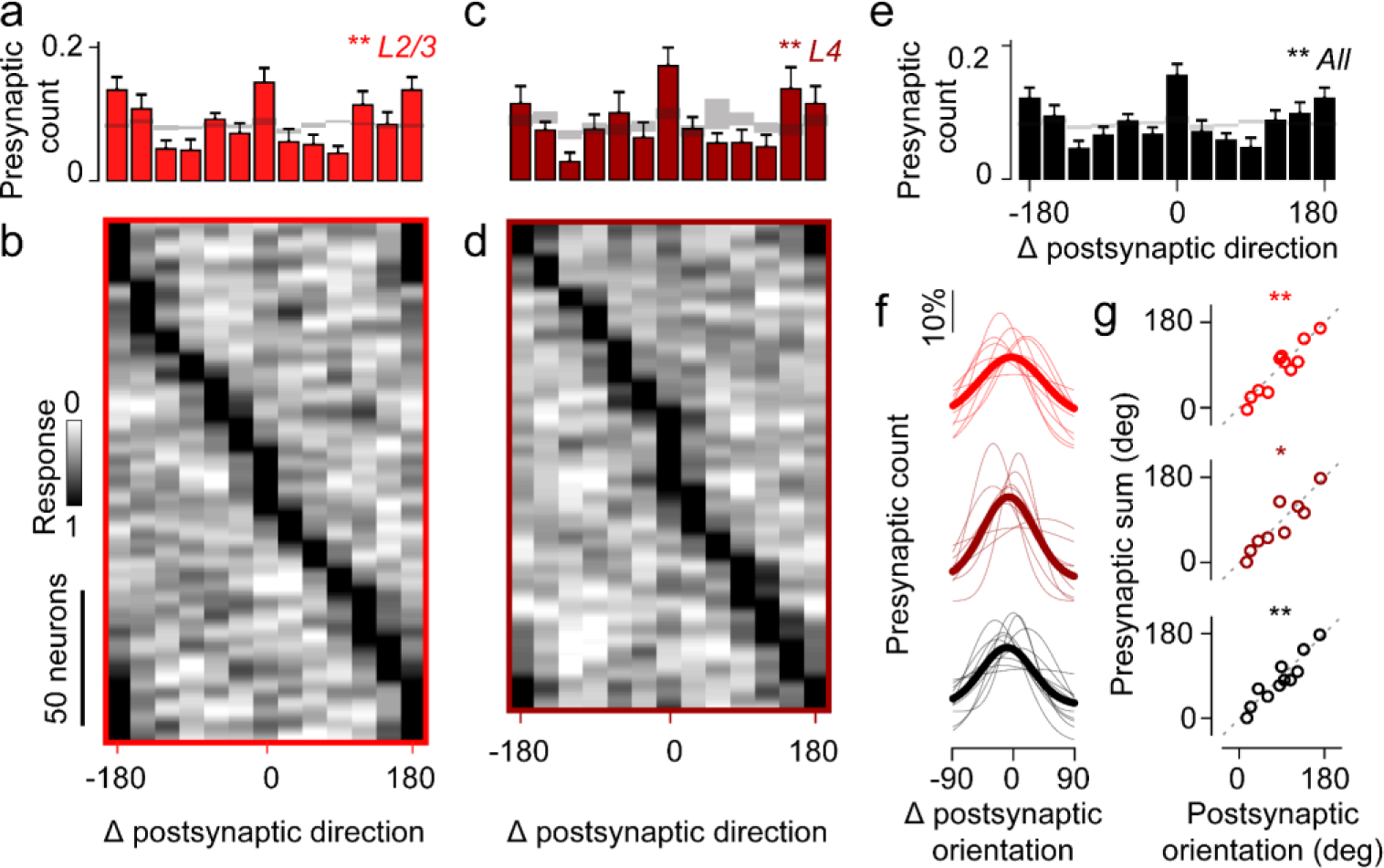
L2/3 neurons receive presynaptic excitation preferentially tuned to their orientation preference. To determine how the tuning of excitatory inputs from different layers contributed to postsynaptic visual selectivity, we presented drifting gratings to the awake mice and recorded from the volume surrounding the postsynaptic neuron, from the surface to superficial L5. (**a**) Distribution of differences in preferred direction between the postsynaptic neurons and their presynaptic ensembles in L2/3 (n=11). Graphs show mean ± s.e. across experiments. Grey shaded area represents confidence intervals calculated from null distributions drawn from the surrounding population. (**b**) Tuning of all excitatory presynaptic neurons in L2/3, normalized and aligned to the preferred direction of the postsynaptic cell. Each row shows the responses of one neuron. (**c**) Same format as **a**, showing the distribution of differences in preferred direction between the postsynaptic neurons and their presynaptic ensembles in L4 (n = 9). (**d**) Same as a **b**, for all the excitatory presynaptic neurons in L4. (**e**) Same format as **a**, showing the distribution of differences in preferred direction between the postsynaptic neurons and their presynaptic ensembles across L2/3, L4 and L5 (n=11). (**f**) Distributions of differences in preferred orientation between the postsynaptic neurons and their presynaptic ensembles in L2/3 (*top,* n=11), in L4 (*middle,* n=9), or pooled across all layers (*bottom,* n=11). Thin curves show fits to individual experiments; the thick curve is fit to the average across experiments (shown in a, c, e). (**g**) The vector sum of these distributions plotted against the postsynaptic orientation preference, for presynaptic ensembles in L2/3 (*top*), in L4 (*middle*), or pooled across layers (*bottom*). The excitatory presynaptic ensemble was significantly aligned to the orientation preference of the postsynaptic neuron (L2/3 r =0.93, p <0.01; L4 r = 0.90, p <0.05; circular correlation). These effects did not result from local biases in neuronal preferences^30,31^, because they were absent in the remaining simultaneously recorded population. Indeed, the tuning of presynaptic ensembles was significantly higher than control surrogate ensembles, matched in size, and crafted from the surrounding population (L2/3 p<0.05 and p<0.01; L4 p<0.05 and p<0.05; for circular correlation and circular V-test). This similarity in preferred orientation, moreover, occasionally extended to preferred direction: 7/11 ensembles had more neurons preferring the postsynaptic direction (5/11 in L2/3; 7/9 in L4). * p<0.05, **p<0.01 circular correlation, circular V-test, or one-way ANOVA.

**Supplementary Figure 5.**
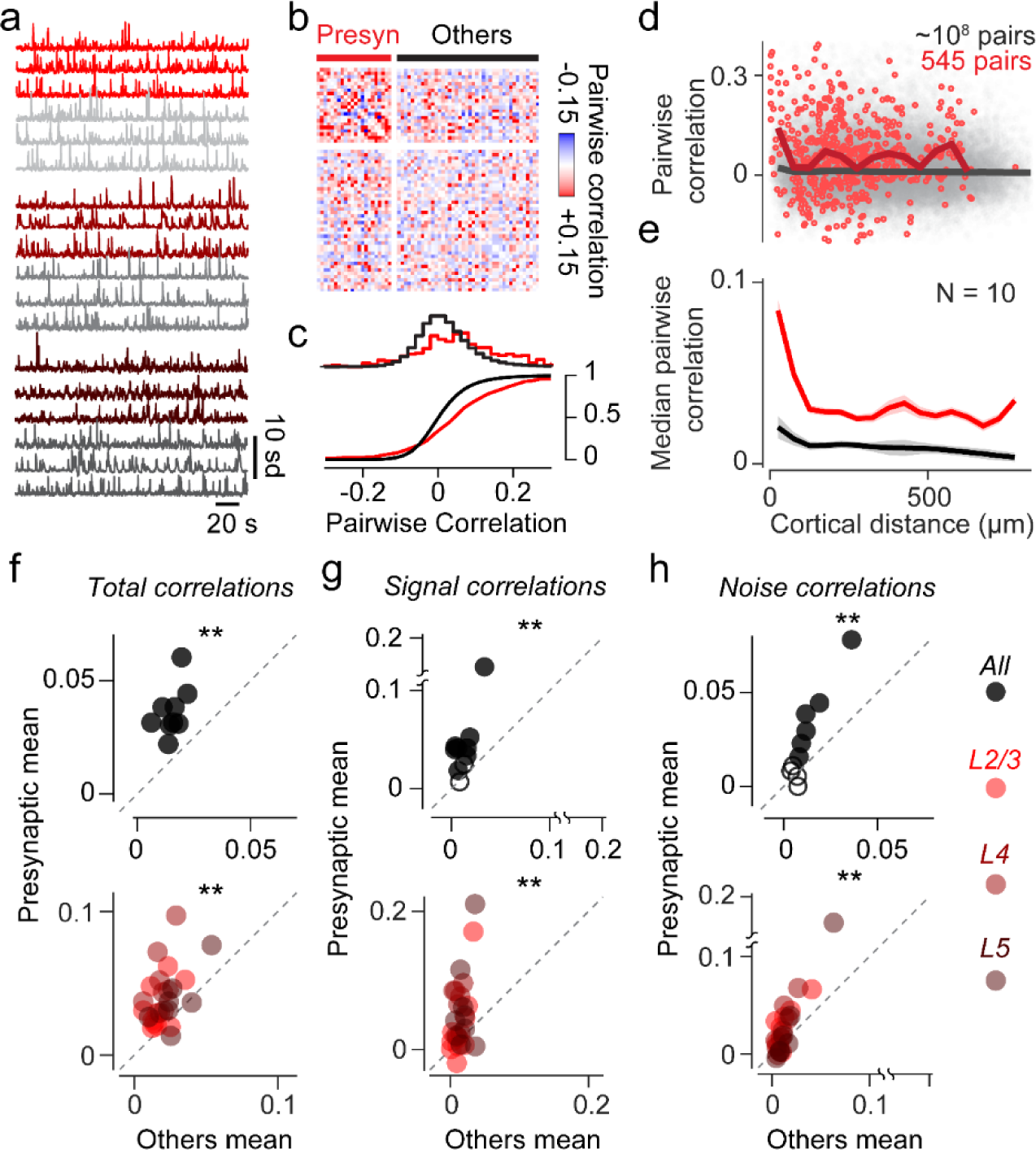
Excitatory presynaptic neurons form networks with higher total, signal and noise correlation. (**a**) Example GCaMP6 fluorescence recordings from L2/3 (*red, bright grey*), L4 (*dark red, grey*), and L5 (*brown, dark grey*) presynaptic neurons (*red*) and simultaneously recorded control neurons (*grey*). (**b**) Pairwise total correlation matrix including 22 presynaptic neurons and 44 randomly chosen control neurons recorded simultaneously. (**c**) Density and cumulative distribution of total pairwise correlations for the presynaptic ensemble and all other control neurons in **b**. (**d**) Total pairwise correlation as a function of inter-somatic lateral distance for pairs of presynaptic neurons (*red*) or control neurons (*grey*). The median dependence of pairwise correlation on inter-somatic distance was computed in 50 µm wide bins (*red and grey lines*). (**e**) Average median distribution of total pairwise correlations as a function of inter-somatic distance for presynaptic neurons (*red*) and the population (*grey*). Data are displayed as mean ± s.e. (N = 10). (**f**) Mean total pairwise correlation across all presynaptic neurons (*top*, N = 10) and for presynaptic neurons grouped by layer (*bottom*, N = 10). Presynaptic ensembles with average pairwise correlation significantly higher than chance are represented by filled circles (*top*) (**g**) Same as **e**, for pairwise signal correlations. (**h**) Same as **e**, for pairwise noise correlations. ** indicates p <0.01, Wilcoxon rank-signed test.

**Supplementary Figure 6.**
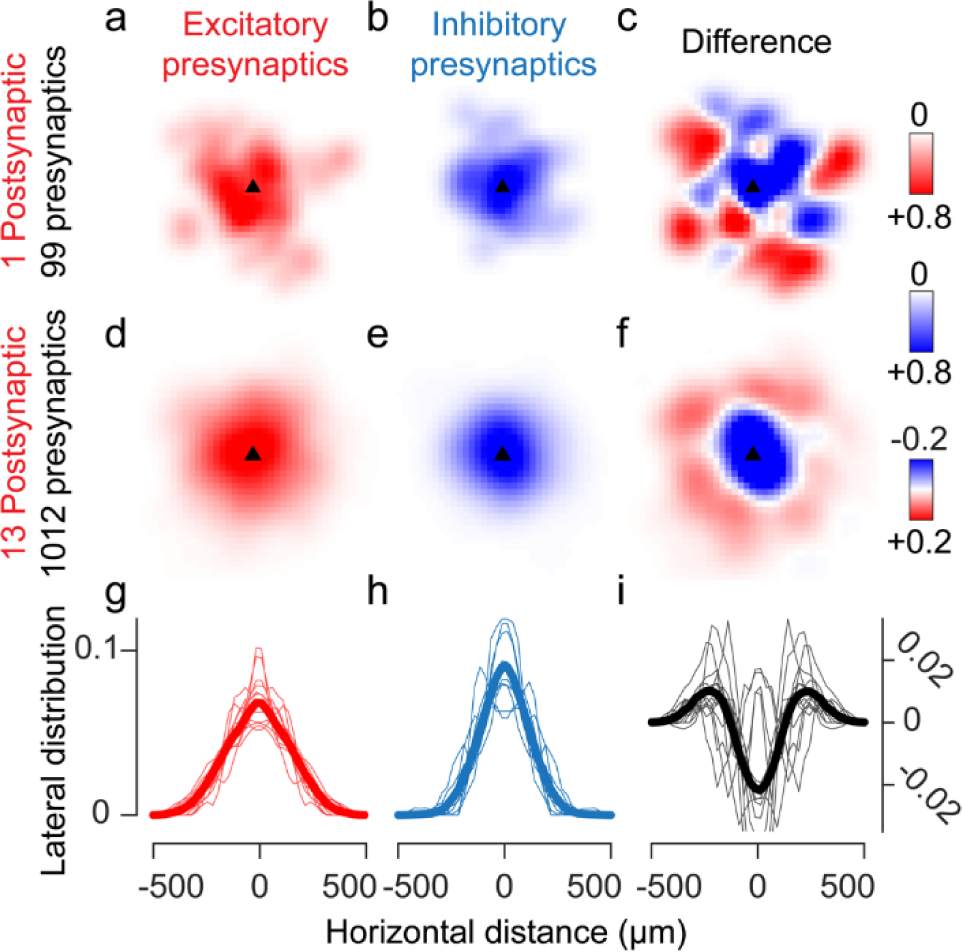
Horizontal distribution of excitatory and inhibitory presynaptic ensembles. (**a**) Horizontal density of excitatory presynaptic neurons around the postsynaptic neuron, for the experiment shown in Figure1c. Density was normalised to its maximum for display purposes. (**a**) Same as **b**, for inhibitory presynaptic neurons. (**c**) Difference between excitatory and inhibitory densities shown in **a** and **b**. (**d-f**) Same as **a-c** for data pooled across experiments (n=13). (**g-i**) Radial distributions for the probability densities in **a-f**, showing individual experiments (*thin lines*), and pooled data (*thick line*). Same horizontal scale as in **a-f**.

**Supplementary Figure 7.**
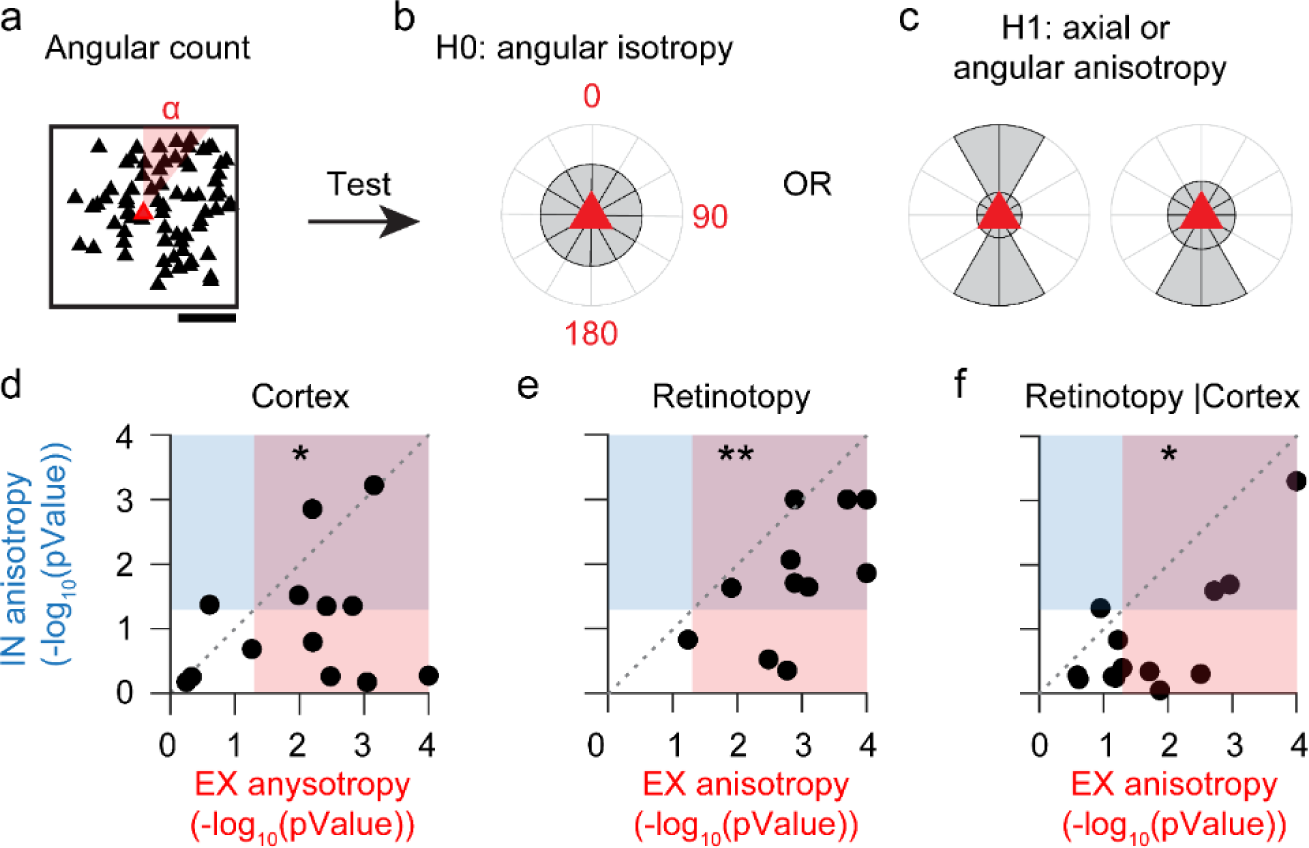
Cortical and retinotopic elongation of excitatory and inhibitory presynaptic distributions. (**a**) Example cortical distribution of presynaptic neurons (*black*) around the postsynaptic neuron (*red*). The position of each neuron can be represented in polar coordinates, where α is the angular coordinate. (**b**) The null hypotheses (*H0*) of angular isotropy: the angular density of presynaptic neurons is uniform around the postsynaptic cell and has high circular variance, signature of equal counts of neurons in each angular sector. (**c**) The alternative hypotheses (*H1*) of angular anisotropy around the postsynaptic cell: axial anisotropy results from pairs of opposite angular sectors containing most of the neurons, resulting in an axially elongated distribution around the postsynaptic cell with low axial variance; angular anisotropy results from angular sectors on one side containing most of the neurons, resulting in an angularly elongated distribution around the postsynaptic cell with low circular variance. The axial and circular variances were computed for each ensemble of presynaptic neuron and were compared against null distributions generated from surrogate, angularly isotropic ensembles. The variance that yielded the largest log_10_(p-value) was used as a measure of anisotropy. (**d**) The cortical anisotropy of the excitatory presynaptic ensembles is plotted against the cortical anisotropy of the inhibitory presynaptic ensembles. (**e**) Same as **d**, for the visual anisotropy. (**f**) Same as in **e**, after correction for biases due to the inhomogeneity and varying magnification factors of the retinotopic map. The correction was achieved testing the data against null distributions designed to be isotropic in cortical space and matched in scale to the data.

